# Hsc70 Ameliorates the Vesicle Recycling Defects Caused by Excess α-Synuclein at Synapses

**DOI:** 10.1101/517524

**Authors:** Susan M. L. Banks, Audrey T. Medeiros, Molly McQuillan, David J. Busch, Ana Sofia Ibarraran-Viniegra, Subhojit Roy, Rui Sousa, Eileen M. Lafer, Jennifer R. Morgan

## Abstract

α-Synuclein overexpression and aggregation are linked to Parkinson’s disease (PD), dementia with Lewy bodies (DLB), and several other neurodegenerative disorders. In addition to effects in the cell body, α-synuclein accumulation occurs at presynapses where the protein is normally localized. While it is generally agreed that excess α-synuclein impairs synaptic vesicle trafficking, the underlying mechanisms are unknown. We show here that acute introduction of excess human α-synuclein at a classic vertebrate synapse, the lamprey reticulospinal synapse, selectively impaired the uncoating of clathrin-coated vesicles (CCVs) during synaptic vesicle recycling, leading to a severe depletion of synaptic vesicles. Furthermore, human α-synuclein and lamprey γ-synuclein both interact *in vitro* with Hsc70, the chaperone protein that uncoats CCVs at synapses. After introducing excess α-synuclein to lamprey axons, Hsc70 availability was reduced at the synapses, suggesting Hsc70 sequestration as a possible mechanism underlying the synaptic vesicle trafficking defects. In support of this hypothesis, increasing the levels of exogenous Hsc70 together with α-synuclein ameliorated the CCV uncoating and vesicle recycling defects. These experiments identify a reduction in Hsc70 availability at synapses, and consequently its function, as the mechanism by which α-synuclein induces synaptic vesicle recycling defects. To our knowledge, this is the first report of a viable chaperone-based strategy for reversing the toxic impacts of excess α-synuclein at synapses, which may be of value for ameliorating synaptic defects in PD and other synuclein-linked diseases.

**SIGNIFICANCE STATEMENT:** Synaptic defects caused by α-synuclein overexpression are linked to cognitive deficits in PD and other diseases. However, the mechanisms by which excess α-synuclein impairs synaptic vesicle trafficking are unknown. Data presented here demonstrate that acute introduction of excess α-synuclein at a classical vertebrate synapse selectively inhibits CCV uncoating, leading to impaired vesicle recycling. Furthermore, increasing α-synuclein reduced synaptic levels of Hsc70, the clathrin uncoating ATPase. Subsequently increasing Hsc70 restored CCV uncoating and improved vesicle recycling. This study identifies a novel molecular mechanism underlying the α-synuclein-induced synaptic defects and presents one viable strategy for reversing them.

## INTRODUCTION

α-Synuclein is a presynaptic protein whose aberrant aggregation causes neurodegeneration in Parkinson’s disease (PD), dementia with Lewy bodies (DLB), and several variants of Alzheimer’s disease (Singleton et al., 2003; Lee and Trojanowski, 2006; Cookson and van der Brug, 2008; Ingelsson, 2016). Inherited forms of PD are linked to multiplication of the α-synuclein gene (*SNCA*), as well as several identified point mutations (e.g. A30P, E46K, A53T), which result in atypical aggregation of α-synuclein protein throughout neurons (Singleton et al., 2003; Lee and Trojanowski, 2006). α-Synuclein aggregation causes synaptic dysfunction, mitochondrial damage, and axonal transport deficits, among other cellular defects (Ingelsson, 2016). Though the mechanisms of α-synuclein toxicity are beginning to emerge, particularly with respect to events in the somata, much less is known about how increased α-synuclein levels affect synapses where the protein normally functions.

Under physiological conditions, α-synuclein functions during synaptic vesicle trafficking via multiple established roles in vesicle clustering/reclustering, SNARE-complex assembly, dilation of the fusion pore, and vesicle recycling (Cabin et al., 2002; Burre et al., 2010; Greten-Harrison et al., 2010; Nemani et al., 2010; Bendor et al., 2013; Vargas et al., 2014; Wang et al., 2014; Logan et al., 2017). However, at present, much less is known about the impacts of excess α-synuclein at synapses, which occurs during disease pathologies. Acutely increasing α-synuclein at several vertebrate synapses leads to an inhibition of synaptic vesicle recycling (Busch et al., 2014; Xu et al., 2016; Eguchi et al., 2017). Overexpression of α-synuclein in animal models of PD induces the formation of small oligomers, microaggregates, and filaments at synapses, following a similar pathological cascade that occurs in other neuronal compartments (Scott et al., 2010; Volpicelli-Daley et al., 2011; Spinelli et al., 2014). This leads to a loss of several critical presynaptic proteins within boutons, which was also observed in the brains of DLB patients (Scott et al., 2010). Perhaps as a consequence, synaptic aggregation of α-synuclein is highly correlated with cognitive deficits in some DLB and PD patients (Kramer and Schulz-Schaeffer, 2007; Schulz-Schaeffer, 2010; Ingelsson, 2016). Despite these indications that excess and/or aggregated α-synuclein negatively impacts presynaptic functions, the underlying mechanisms are unknown.

Lamprey giant reticulospinal (RS) synapses provide an excellent model for assessing how excess α-synuclein affects vertebrate synapses. RS synapses are ideal for these studies because they are amenable to acute perturbations of presynaptic processes (Brodin and Shupliakov, 2006; Walsh et al., 2018), thus permitting a direct evaluation of the effects of excess α-synuclein without inducing molecular compensation that occurs after overexpression (Nemani et al., 2010; Scott et al., 2010). Furthermore, the large size of vesicle clusters at RS synapses facilitates detailed ultrastructural analyses of synaptic vesicle trafficking events, allowing us to determine the underlying mechanisms (Morgan et al., 2004; Morgan et al., 2013; Busch et al., 2014). We previously reported that acute introduction of excess human α-synuclein inhibited synaptic vesicle recycling at RS synapses, including effects on clathrin-mediated endocytosis (Busch et al., 2014; Medeiros et al., 2017; Medeiros et al., 2018). Similarly, α-synuclein also inhibited vesicle endocytosis, but not exocytosis, at mammalian synapses (Xu et al., 2016; Eguchi et al., 2017).

We report here the mechanism by which excess α-synuclein induces synaptic vesicle recycling defects and present a novel strategy for ameliorating them. When introduced acutely, α-synuclein selectively impaired the uncoating of clathrin-coated vesicles (CCVs) during synaptic vesicle recycling, leading to a depletion of the vesicle cluster. We further show that human α-synuclein, lamprey γ-synuclein, and several PD-linked mutants directly associate *in vitro* with Hsc70, the chaperone protein that uncoats CCVs at synapses, thus identifying an interaction that may affect synapses *in vivo.* Indeed, excess α-synuclein reduced Hsc70 availability at stimulated synapses, suggesting Hsc70 sequestration as a possible mechanism underlying the synaptic defects. Consequently, co-injection of exogenous Hsc70 together with α-synuclein largely ameliorated the synaptic vesicle trafficking defects. Thus, Hsc70 is an *in vivo* target of excess α-synuclein at synapses, and increasing Hsc70 function reverses the deleterious impacts.

## MATERIALS AND METHODS

### Recombinant proteins

Cloning of recombinant GST-tagged human α-synuclein and His-tagged bovine Hsc70 used for biochemistry experiments was as described in (Wilbanks et al., 1995; Busch and Morgan, 2012; Busch et al., 2014; Sousa et al., 2016). Recombinant proteins were expressed in BL21-CodonPlus (DE3)-RILP Competent Cells (Agilent Technologies, Santa Clara, CA) and purified using Glutathione Sepharose 4B Media (GE Healthcare; Pittsburgh, PA) or Ni-NTA resin (Thermo Fisher Scientific; Waltham, MA). Untagged human α-synuclein used in the microinjection experiments was obtained from rPeptide (Bogart, GA).

### Acute perturbations and electron microscopy

All animal procedures were approved by the Institutional Animal Care and Use Committee at the MBL in accordance with standards set by the National Institutes of Health. Lampreys (*Petromyzon marinus;* 11-13 cm; 5-7 years old M/F) were anesthetized in 0.1 g/L MS-222 (Western Chemical Inc.; Ferndale, WA). Spinal cord pieces (2-3 cm) were dissected and pinned ventral side up in a Sylgard lined dish. Axonal microinjections were performed as described in Walsh et al., 2018. First, human α-synuclein was diluted in lamprey internal solution (180 mM KCl, 10 mM HEPES K+, pH 7.4) to a pipet concentration of 130-160 μM. In some experiments, an equimolar concentration of recombinant bovine Hsc70 was included in the solution. Proteins were then loaded into glass microelectrodes (20-25 MΩ) and microinjected into giant RS axons using small pulses of N_2_ (5-20 ms, 30-50 psi, 0.2 Hz) delivered through a picospritzer. Fluorescein (10 kDa) or tetramethylrhodamine (70 kDa) dextrans (100 μM; Thermo Fisher; Waltham, MA), approximating the molecular weights of α-synuclein and Hsc70, respectively, were included in the pipets and co-injected in order to visualize the proteins’ diffusion rates in the axons. Axonal injections resulted in a 10-20x dilution of the proteins. Thus, the final axonal concentrations of α-synuclein and Hsc70 were ~7-16 μM. Axons were subsequently stimulated with action potentials (20 Hz, 5 min) using current injections (30-60 nA; 1 ms) to induce synaptic vesicle exocytosis/endocytosis.

Spinal cords were fixed immediately after stimulation (3% glutaraldehyde, 2% paraformaldehyde in 0.1 M Na cacodylate, pH 7.4), processed for EM, sectioned at 70 nm, and counterstained with uranyl acetate and lead citrate, as described in (Morgan et al., 2013; Busch et al., 2014; Walsh et al., 2018). Images were obtained at 37,000x or 59,000x magnification (n=22-30 synapses, 2 lampreys/condition) using a JEOL JEM 200CX electron microscope. Images were collected at distances surrounding the injection site (20-150 μm) where the protein concentration was measurable based on the diffusion of the co-injected fluorescent dye (i.e. the experimental condition), as well as distances farther from the injection site (150-700 μm) where no protein had diffused (i.e. the controls). 3D reconstructions were generated from 5 serial electron micrographs using Reconstruct software (Fiala, 2005), as described in (Busch et al., 2014; Medeiros et al., 2017). One image per synapse, taken at or near the center of the active zone, was selected for morphometric analysis. A researcher blinded to the experimental conditions performed the morphometric analyses on all synaptic membranes within a 1-μm radius of the active zone using FIJI 2.0.0 (Morgan et al., 2004; Morgan et al., 2013; Busch et al., 2014). These included synaptic vesicles, plasma membrane, cisternae, and clathrin-coated pits and vesicles. Synaptic vesicles were defined as small, clear round vesicles <100 nm in diameter, while “cisternae” were defined as larger vesicles that were >100nm in diameter, as in our previous studies (Busch et al., 2014; Medeiros et al., 2017). Plasma membrane evaginations were determined by drawing a straight line from the edge of the active zone to the nearest position on the axolemma, on both sides of the synapse, and then measuring the curved distance between these points; the mean value per synapse was recorded. Clathrin-coated pits and vesicles were staged as described in (Morgan et al., 2004). After obtaining each membrane measurement, a total membrane analysis was performed on each synapse in order to determine how membrane was redistributed with each perturbation. Here, SV and CCP/V membrane areas were calculated by multiplying the surface area of a sphere (4πr^2^) by the number of each type of vesicle at each synapse. Plasma membrane and cisternae areas were obtained by multiplying the length of membrane evaginations and summed cisternae perimeters, respectively, by the section thickness (70 nm). Graphing and statistical analyses, including Student’s t-Tests and ANOVA, were performed in Origin 7.0 (OriginLab Corp; Northampton, MA). Data were reported as the mean value per section per synapse.

### GSTpull-downs

Protein lysates from rat brains or lamprey CNS (brains and spinal cords) were prepared in HKET buffer (25 mM HEPES K+, 150 mM KCl, 1 mM EDTA, 1% Triton X-100, pH 7.4) containing protease inhibitors. GST pull downs were performed using 50-100 μg of GST-tagged proteins and either 1-5 mg brain extracts or 100 μg purified recombinant proteins. After performing the pull downs, the bound proteins were run on 10% SDS-PAGE gels and transferred to nitrocellulose membranes. For Western blotting, antibodies were diluted in TBST (20 mM Tris pH 7.6, 150 mM NaCl, 0.1% Tween 20) with 1% dry milk or 5% BSA. Primary antibodies were used at 1:1000 and included: mouse monoclonal anti-β1/β2-adaptins (clone 100/1; Sigma, St. Louis, MO); mouse monoclonal anti-clathrin heavy chain (clone 23; BD Biosciences, San Jose, CA); mouse monoclonal anti-dynamin (clone 41; BD Biosciences, San Jose, CA); mouse monoclonal anti-synaptojanin-1 (clone 26; BD Biosciences, San Jose, CA); mouse monoclonal anti-auxilin (gift from Ernst Ungewickell) (Ungewickell et al., 1995); rabbit polyclonal anti-Hsc70 (ARP48445; Aviva Systems Biology, San Diego, CA); rabbit polyclonal anti-Hsc70 (SPA-816; Enzo Life Sciences, Farmingdale, NY). Secondary antibodies were used at 1:1000-1:4000 and included HRP-conjugated goat anti-rabbit, anti-mouse, and anti-rat IgGs (H+L) (Thermo Scientific, Waltham, MA). Protein bands were detected using Pierce^™^ ECL Western blotting substrate (Thermo Scientific, Waltham, MA). Band intensity was measured using FIJI 2.0.0 software. Data shown in graphs indicate mean ± SEM from n=3-5 experiments.

### Clathrin uncoating assays

Clathrin cages were assembled with 1 μM recombinant bovine brain clathrin and 0.1 μM auxilin, as described in Sousa et al., 2016. CCVs were freshly purified from bovine brains as described in (Keen et al., 1979; Nandi et al., 1982). To visualize the clathrin cages and CCVs, freshly glow-discharged copper grids (EM Sciences; Hatfield, PM) were floated onto a drop of each sample for 5 minutes, followed by six washes in distilled H2O, counterstaining in 1% uranyl acetate for 3 minutes in the dark. After drying, the grids were imaged on a JEOL JEM 200CX electron microscope at 100 kV using 100,000x magnification. Clathrin disassembly from clathrin cages and purified CCVs was measured *in vitro* using the dynamic light scattering assay described in (Schuermann et al., 2008; Morgan et al., 2013). Briefly, clathrin cages or CCVs were incubated in 20 mM imidazole, 25 mM KCl, 10 mM (NH4)2SO4, 2 mM Mg acetate, and 2 mM DTT, pH 7.0, at 25°C. Clathrin disassembly was initiated by adding 2 μM bovine Hsc70 preincubated with 1 mM ATP with or without 10 μM recombinant human α-synuclein (rPeptide), and monitored by dynamic light scattering using a Wyatt DynaPro. Data were plotted using Origin 7.0 software.

### Immunofluorescence (IF) at lamprey synapses

Recombinant human α-synuclein was microinjected into giant RS axons as described above for the EM experiments, after which spinal cords were stimulated using high K+ (50 mM) Ringer for 10 min (Wickelgren et al., 1985). In some experiments (Fig. 5G-H), spinal cords were stimulated using action potentials (20 Hz, 5 min). Following stimulation, spinal cords were fixed in 4% paraformaldehyde in 0.1 M PBS, pH 7.4, for 3 hrs, washed in 0.1 M PBS, and incubated for 1 hr in blocking buffer (10% Normal Goat Serum; Thermo Fisher Scientific; Waltham, MA) containing 0.3% Triton X-100. Primary antibody incubations were overnight at 4°C, followed by 5 x 1 hour washes in wash buffer (20 mM Na phosphate buffer, 450 mM NaCl, 0.3% Triton X-100, pH 7.4). Primary antibodies included: mouse monoclonal anti-SV2 antibody, which was deposited to DSHB by K.M. Buckley (1:100; DSHB, Iowa City, IA) (Buckley and Kelly, 1985); rabbit polyclonal anti-Hsc70 (ARP48445; Aviva Systems Biology, Corp., San Diego, CA); and rat monoclonal anti-Hsc70 (SPA-815; Enzo Life Sciences, Farmingdale, NY). Secondary antibodies used were Alexa Fluor^®^ 594 goat anti-mouse IgG (H+L) (1:200); Alexa Fluor^®^ 488 goat anti-rabbit IgG (H+L) (ThermoFisher; Waltham, MA); or DyLight^®^ 488 goat anti-rat IgG (H+L) (1:100) (Thermo Fisher; Waltham, MA). After immunostaining, synapses were imaged within intact whole mounted spinal cords using a Zeiss LSM510 Meta confocal on an Axioskop 2FS microscope. Images were acquired using a Zeiss 40x, 0.8 NA Achroplan objective with 3x optical zoom. All analyses on synapses were performed in FIJI 2.0.0 as follows. Giant RS synapses were first identified using the SV2 labeling, and then the associated Hsc70 puncta (≥2x background intensity) were identified and measured within a 1-μm radius of the synapses, representing the periactive zone where endocytosis occurs (see Fig. 5A). The percentage of synapses containing Hsc70 puncta within each axon, as well as the average number of Hsc70 puncta per synapse, was calculated. All graphing and statistical analyses were performed in Prism 8.0.0 software, including outlier and normality tests (Shapiro-Wilk; D’Agostino & Pearson), as well as statistical comparisons using Student’s t-Test or ANOVA as required (GraphPad Software; San Diego, CA).

## RESULTS

### Excess α-synuclein impairs clathrin uncoating *in vivo* during synaptic vesicle recycling

In order to determine the possible cellular and molecular targets of α-synuclein, we first needed to further determine which stage or stages of clathrin-mediated endocytosis (CME) were preferentially affected. Briefly, clathrin-mediated vesicle recycling is initiated when clathrin adaptors, AP180 and AP2, recruit clathrin triskelia to the plasma membrane, promoting their assembly (Fig. 1A) (Morgan et al., 1999; Saheki and De Camilli, 2012). After maturation of the clathrin-coated pit (CCP), vesicle fission occurs through the actions of dynamin, a large GTPase that is abundant at synapses (Takei et al., 1995; Antonny et al., 2016). Synaptojanin is recruited to the neck of the CCP to assist during fission and subsequently in clathrin uncoating (Cremona et al., 1999; Chang-Ileto et al., 2011). After vesicle fission is complete, the free CCVs are uncoated by the actions of the chaperone protein Hsc70 and its co-chaperone, auxilin (Ungewickell et al., 1995; Morgan et al., 2001). Uncoated vesicles are then refilled with neurotransmitter molecules and returned to the vesicle cluster for subsequent bouts of exocytosis.

**FIGURE 1.**
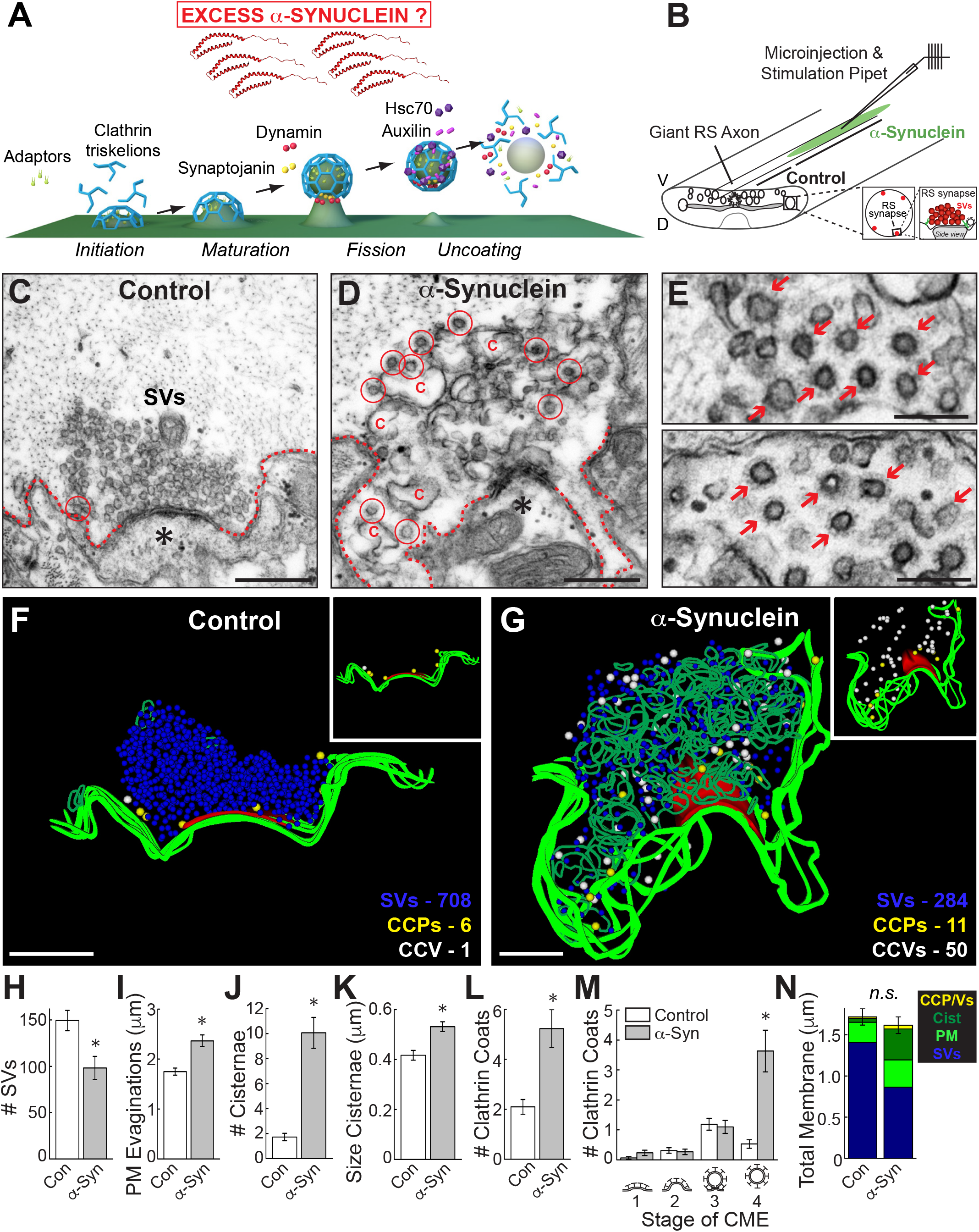
Excess α-synuclein impairs CCV uncoating during synaptic vesicle recycling. **A.** Model showing the major stages of clathrin-mediated synaptic vesicle endocytosis and several molecular players. The goal of the study is to determine how excess α-synuclein affects this process. Graphics generated by Jack Cook (Woods Hole Oceanographic Institution) using Cinema 4D. **B.** Diagram of the lamprey spinal cord showing microinjection strategy and location of reticulospinal (RS) synapses. D=dorsal; V=ventral. **C-D.** Electron micrographs of untreated, control lamprey synapses stimulated at 20 Hz for 5 min. After stimulation, control synapses have large synaptic vesicle (SV) clusters, shallow plasma membrane (PM) evaginations (dotted lines), and only a few CCP/Vs (circles). In contrast, synapses treated with excess human α-synuclein had smaller SV clusters, larger PM evaginations, and greater numbers of CCP/Vs, indicative of a vesicle recycling defect. Asterisks mark postsynaptic spines. Scale bars = 500 nm. **E.** Insets show clusters of free CCVs at synapses after treatment with α-synuclein (arrows). Scale bars = 200 nm. **F-G.** 3D reconstructions comparing control and α-synuclein treated synapses. α-Synuclein caused severe endocytic defects, including a striking increase in CCVs (white spheres) and cisternae (dark green traces). Insets show the distributions of CCPs (yellow spheres) and CCVs (white spheres). While CCP/Vs at control synapses are sparse and localized near the plasma membrane at control synapses (F, inset), α-synuclein treated synapses exhibited large numbers of free CCVs throughout the synaptic area with little change in CCPs (F, inset). Active zone is shown in red. Scale bars = 500 nm. **H-N.** The SV recycling defect induced by α-synuclein is demonstrated by a loss of SVs, which was compensated by larger PM evaginations and greater numbers of cisternae and CCP/Vs. The selective increase in CCVs (stage 4) indicates a clathrin uncoating defect. Bars represent mean ± SEM (per section, per synapse) from n=30-33 synapses, 2 axons/condition. Asterisks denote significance (p<0.05) by Student’s t-Test (H-L, N) or ANOVA (M).

As in our prior studies, we microinjected recombinant monomeric human α-synuclein into lamprey giant axons, thereby delivering the protein directly to presynapses (Fig. 1B). The axons were subsequently stimulated (20 Hz, 5 min), fixed, and processed for standard transmission electron microscopy. After injection, the final axonal concentration of α-synuclein was estimated to be ~7-16 μM, which is ~2-5 times greater than measurements of endogenous α-synuclein at synapses and commensurate with overexpression levels in mammalian PD models and human patients (Add a human ref here) (Nemani et al., 2010; Scott et al., 2010; Westphal and Chandra, 2013). At control synapses, local synaptic vesicle recycling was efficient enough to maintain a large synaptic vesicle cluster, and very few CCP/Vs were observed (Fig. 1C, F). In contrast, after injection of α-synuclein, synapses were dramatically altered due to deficits in synaptic vesicle recycling. Specifically, synapses treated with α-synuclein exhibited fewer synaptic vesicles, expanded plasma membrane evaginations, increased numbers of large atypical “cisternae”, and abundant CCVs (Fig. 1D, G). “Cisternae” were classified as any irregular-shaped vesicles with a diameter >100 nm (Busch et al., 2014; Medeiros et al., 2017). These cisternae may be recycling endosomes or bulk endosomes, because they often had CCPs budding from them and could sometimes be traced back to the plasma membrane, as previously reported (Busch et al., 2014). Strikingly, atypical clusters of free CCVs were observed at synapses treated with excess α-synuclein, a phenotype typically associated with defective clathrin uncoating (Fig. 1E, arrows) (Cremona et al., 1999; Morgan et al., 2001). 3D reconstructions generated from serial micrographs revealed the gross alterations in synaptic structure caused by α-synuclein (Fig. 1F-G). Whereas control synapses exhibited only a few CCPs and CCVs close to the plasma membrane, the α-synuclein treated synapses exhibited dozens of CCVs that were dispersed throughout the synaptic area (Fig. 1F-G, insets).

We performed a quantitative morphometric analysis of all synaptic membranes within 1 micron of the active zone and confirmed that excess α–synuclein impaired synaptic vesicle recycling, as demonstrated by a loss of synaptic vesicles that was compensated by larger plasma membrane evaginations (Fig. 1H-I) (*Synaptic Vesicles* - Control: 150 ± 11 SVs/section, n=31 synapses, 2 axons; α-Synuclein: 98 ± 12 SVs; n=30 synapses, 2 axons; Student’s t-Test; p<0.005) (*Plasma Membrane* – Control: 1.75 ± 0.07 μm, n=32 synapses, 2 axons; α-Synuclein: 2.37 ± 0.12 μm; n=30 synapses, 2 axons; Student’s t-Test; p<0.00005). The remaining synaptic vesicles were of normal size (diameter) (Control: 54.7 ± 0.6 nm, n = 200 SVs, 10 synapses; α-Synuclein: 52.9 ± 0.9 nm; n=200 SVs, 10 synapses; Student’s t-Test; p=0.10). Corroborating the vesicle recycling defect, the number and the size (perimeter) of the cisternae were also significantly increased (Fig. 1J-K) (*# Cisternae* – Control: 1.7 ± 0.3 cisternae, n=32 synapses, 2 axons; α-Synuclein: 10.1 ± 1.2 cisternae; n=30 synapses, 2 axons; Student’s t-Test; p<0.00001) (*Size Cisternae* – Control: 0.42 ± 0.02 μm, n=55 cisternae, 32 synapses; α-Synuclein: 0.53 ± 0.02 μm; n=302 cisternae, 30 synapses; Student’s t-test; p<0.05). Supporting effects on CME, the total numbers of clathrin-coated pits and vesicles (CCPs+CCVs) were increased nearly 3-fold (Fig. 1L) (*# Clathrin coats* – Control: 2.1 ± 0.3 coats, n=32 synapses, 2 axons; α-Synuclein: 5.2 ± 0.8 coats, n=30 synapses, 2 axons; Student’s t-Test; p<0.0005). Further dissection of the stages of CME revealed that α-synuclein selectively increased the number of free CCVs (stage 4) without significantly altering the earlier stages (Fig. 1M) (*Stage 1–* Control: 0.06 ± 0.04 CCPs/section; α-Synuclein: 0.23 ± 0.09 CCPs; *Stage 2–* Control: 0.31 ± 0.09 CCPs, α-Synuclein: 0.27 ± 0.10 CCPs; *Stage 3-* Control: 1.19 ± 0.20 CCPs; α-Syn: 1.10 ± 0.22 CCPs; *Stage 4-* Control: 0.53 ± 0.14 CCVs, α-Synuclein: 3.63 ± 0.70 CCVs; n=30-32 synapses, 2 axons; ANOVA p<0.0000005, Tukey’s post hoc). A total membrane analysis shows clearly that the loss of synaptic vesicle membrane area was compensated by an expansion of the plasma membrane, cisternae, and CCP/Vs (Fig. 1N) (Control 1.7 ± 0.1 μm^2^; α-Syn: 1.6 ± 0.1 μm^2^; n=30-32 synapses; Student’s t-Test; p=0.47). Taken together, these data indicate that excess α-synuclein impairs clathrin-mediated synaptic vesicle recycling with selective effects on CCV uncoating (Medeiros et al., 2017, 2018).

### α-Synuclein interacts with Hsc70, the uncoating ATPase at synapses

To determine how α-synuclein impairs clathrin uncoating, we next tested for possible interactions between α-synuclein and several key players that mediate CME at synapses, including those involved in the uncoating process (Fig. 1A) (Saheki and De Camilli, 2012). Human α-synuclein consists of a highly conserved N-terminal domain (NTD; a.a. 1-95), which folds into an amphipathic α-helix upon interaction with small vesicles and includes the non-Aβ component (NAC) domain (a.a. 61-95), followed by a less-structured acidic C-terminal domain (a.a. 96-140) (Fig. 2A). GST-tagged human α-synuclein was used in pull-down experiments to test for any binding partners isolated from rat brain protein extracts. In the GST pull-downs, no interactions were observed between α-synuclein and β-adaptin (an AP2 subunit), clathrin heavy chain, dynamin, synaptojanin, or auxilin (Fig. 2B). In contrast, full-length α-synuclein and its highly conserved NTD, selectively pulled down Hsc70, the chaperone protein that uncoats clathrin at synapses via its ATPase activity (Fig. 2C, top). Similarly, GST-α-synuclein and NTD pulled down Hsc70 from protein extracts isolated from the lamprey brain and spinal cord tissues, demonstrating conservation of the interaction (Fig. 2C, bottom). We repeated the pull-downs using GST-tagged lamprey γ-synuclein, which is the most highly expressed isoform in the lamprey giant RS neurons (Busch and Morgan, 2012). Full-length lamprey γ-synuclein is 56% identical and 63% similar to human α-synuclein, and its α-helical NTD is 67% identical and 90% similar to the corresponding region of human α-synuclein (Fig. 2A) (Busch and Morgan, 2012; Busch et al., 2014). Further corroborating this conserved interaction, lamprey γ-synuclein and its NTD also pulled down Hsc70 from rat brain extracts (Fig. 2D).

**FIGURE 2.**
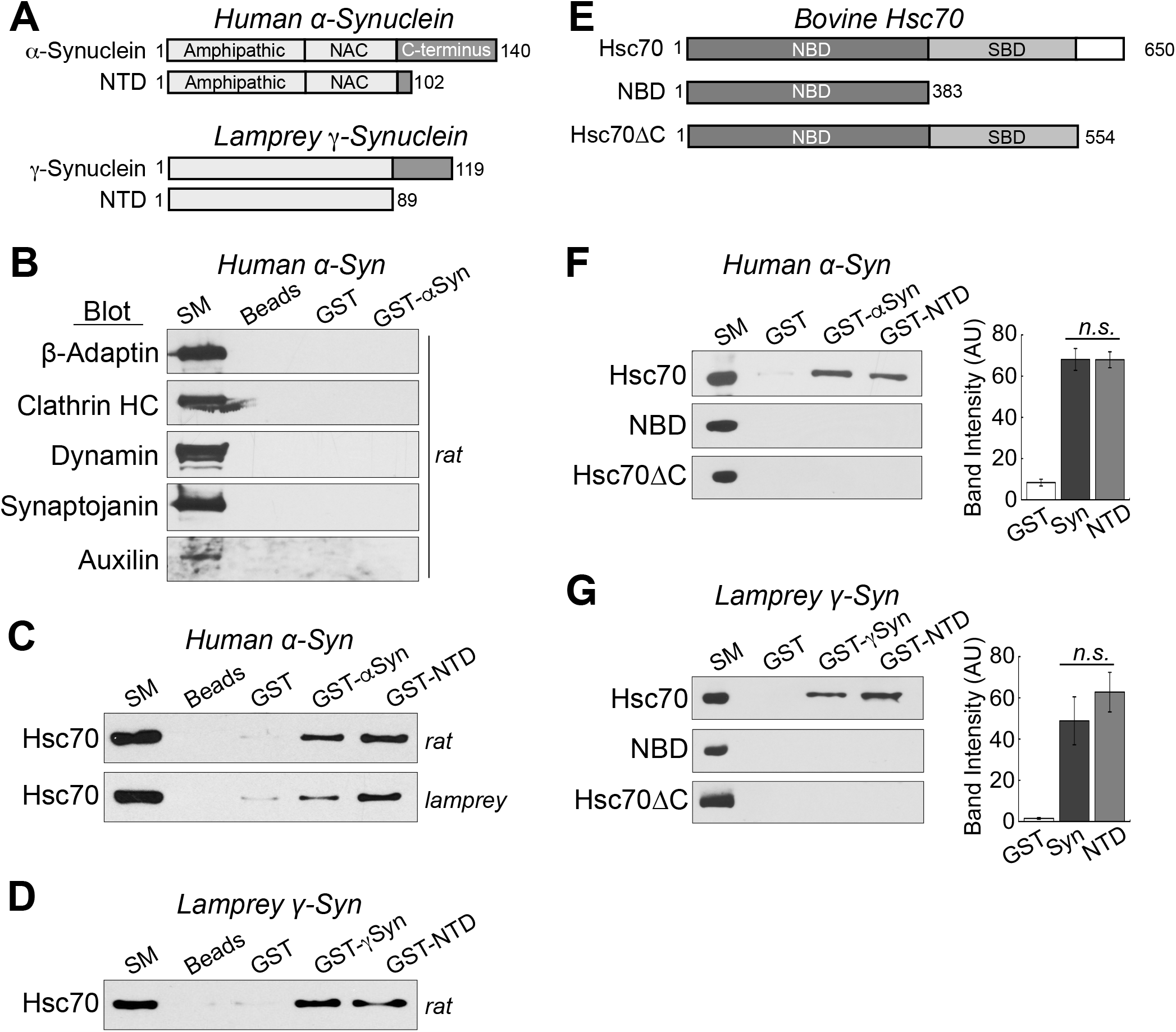
α-Synuclein interacts directly with Hsc70, the chaperone protein that uncoats CCVs during vesicle recycling. **A.** Domain diagrams of full-length human α-synuclein, lamprey γ-synuclein and their respective N-terminal domains (NTDs). NAC = non-amyloid component. **B.** GST pull downs from rat brain lysates revealed no detectable interactions between α-synuclein and several major components of clathrin-mediated endocytosis including AP2, clathrin, dynamin, synaptojanin, and auxilin. SM=starting material. Blots shown are representative of n=3 experiments. **C.** In contrast, human α-synuclein and its NTD pulled down Hsc70 from rat brain and lamprey CNS lysates. **D.** Similarly, lamprey γ-synuclein pulled down Hsc70 from rat brain lysates, demonstrating conservation of the interaction. **E.** Domain diagrams of bovine Hsc70 and several truncations used in the experiments. NBD = nucleotide binding domain. SBD = substrate binding domain. **F-G.** In direct binding assays, both human α-synuclein and lamprey γ-synuclein, and their NTDs, pulled down recombinant bovine Hsc70. No interactions were detected with either NBD or Hsc70ΔC, indicating a role for the C-terminus. Bars represent mean ± SEM from n=3 experiments. AU = arbitrary units. *n.s.* = not significant by ANOVA (p>0.05).

To determine whether the Hsc70/α-synuclein interaction is direct, the GST pull-downs were repeated using purified, recombinant bovine Hsc70. Hsc70 consists of a highly conserved N-terminal nucleotide-binding domain (NBD), and a substrate binding domain (SBD) with a more variable 10 kDa C-terminal region (Fig. 2E). While the affinity of Hsc70 for typical client polypeptides is normally modulated by nucleotides (ATP/ADP), we performed the pull-downs without nucleotides because prior studies reported more effective sequestration of α-synuclein by Hsc70 in fibrillation assays under nucleotide-free conditions (Pemberton et al., 2011; Redeker et al., 2012). Under these conditions, GST-tagged human α-synuclein and its NTD pulled down bovine Hsc70, indicating a direct interaction (Fig. 2F, top) (GST: 8.3 ± 1.6 AU; GST-Syn: 68.1 ± 5.3 AU; GST-NTD: 67.9 ± 3.9 AU; n=3; ANOVA p=5 x 10^−5^; Tukey’s post hoc). Similarly, lamprey γ-synuclein and its NTD also pulled down bovine Hsc70 in these direct binding assays (Fig. 2G, top) (GST: 1.5 ± 0.4 AU; GST-Syn: 48.9 ± 11.6 AU; GST-NTD: 62.8 ± 9.6 AU; n=3; ANOVA p=0.006; Tukey’s post hoc). To further map the interaction, we additionally tested the binding of α-synuclein to several Hsc70 truncation mutants. One truncation consisted of only the nucleotide binding domain (NBD; a.a. 1-383), and the other contained the NBD and substrate binding domain but lacked the C-terminal 10 kD region (Hsc70ΔC; a.a. 1-554) (Fig. 2E)(Flaherty et al., 1990; Wilbanks et al., 1995). Neither human α-synuclein, lamprey γ-synuclein, nor the NTDs pulled down the Hsc70 truncation mutants that were missing the C-terminal fragments, indicating a role for the C-terminus in mediating or stabilizing the interaction (Fig. 2F-G, middle and bottom). Taken together, these data indicate that the N-terminus of α-synuclein interacts directly with Hsc70 via an interaction that is conserved between synuclein orthologs and disrupted by the deletion of the C-terminus of Hsc70.

We repeated the pull downs using several point mutants of α-synuclein that are linked to PD: A30P, E46K, and A53T (Fig. 3A). All three of the point mutants bound directly and robustly to bovine Hsc70 (Fig. 3B-D). While A30P and A53T did not exhibit any statistically significant differences in binding efficacy, as compared to wild type α-synuclein, E46K binding to Hsc70 was slightly reduced (Fig. 3B-D) (GST: 1.1 ± 0.3 AU; GST-Syn: 75.0 ± 6.9 AU; GST-A30P: 57.0 ± 11.8 AU; n=5; ANOVA p<0.0001; Tukey’s post hoc) (GST: 11.5 ± 2.3 AU; GST-Syn: 97.8 ± 5.5 AU; GST-E46K: 75.2 ± 6.3 AU; n=3; ANOVA p<0.0001; Tukey’s post hoc) (GST: 6.1 ± 4.4 AU; GST-Syn: 68.0 ± 6.8 AU; GST-A53T: 67.1 ± 3.2 AU; n=5; ANOVA p<0.0001; Tukey’s post hoc).

**FIGURE 3.**
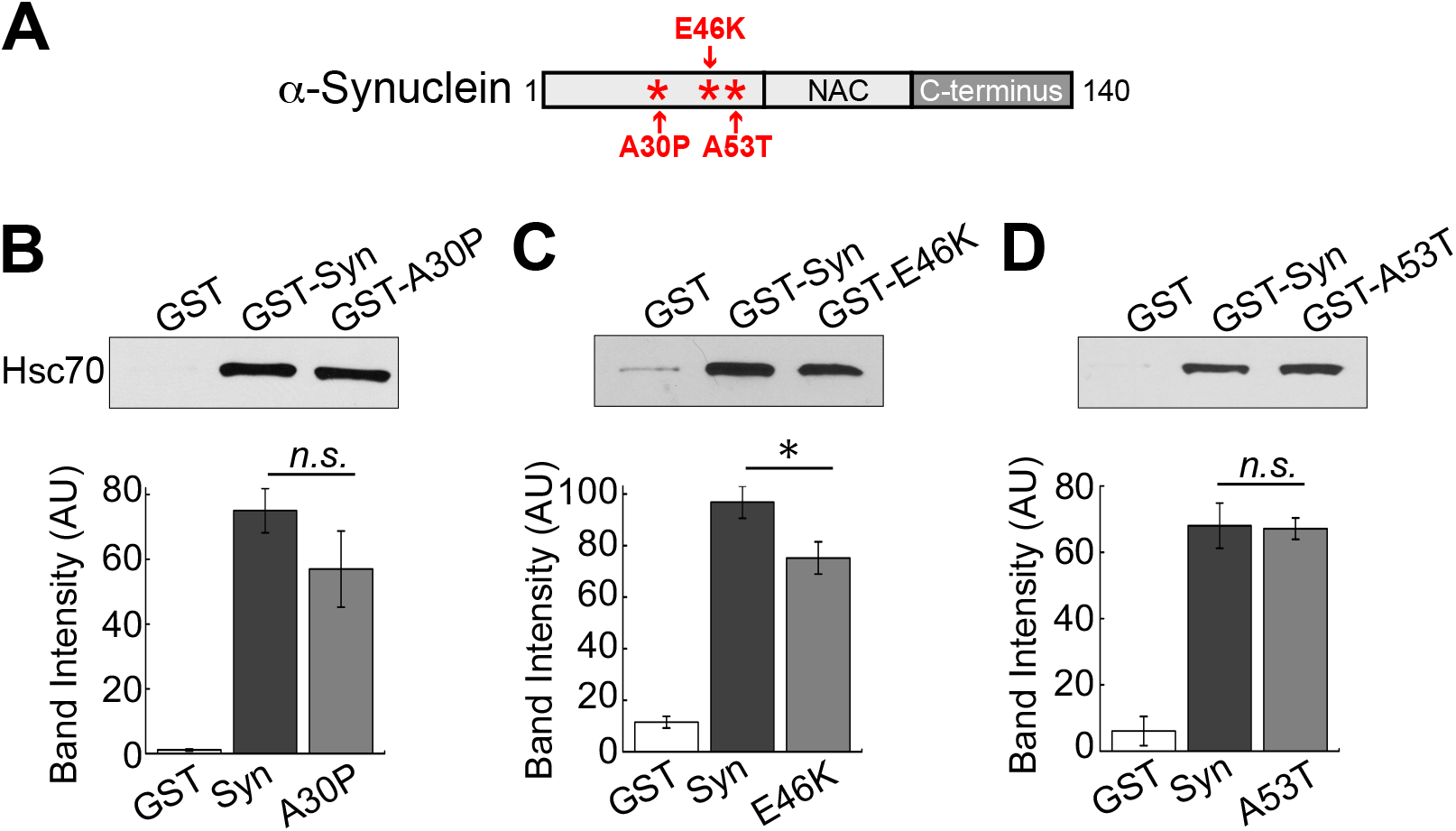
Parkinson’s disease-associated α-synuclein mutants interact with Hsc70. **A.** Diagram showing the locations of PD-linked point mutations A30P, E46K, and A53T, which occur within the helical N-terminal domain of α-synuclein. **B-D.** In direct binding assays, GST-tagged A30P, E46K, and A53T all pulled down Hsc70 to a similar degree as wild type α-synuclein. Only binding to E46K was slightly reduced. Bars represent mean ± SEM from n=3-5 experiments. AU = arbitrary units. Asterisk indicates significance (p=0.04), and n.s. = not significant by ANOVA.

### α-Synuclein does not affect Hsc70-mediated clathrin disassembly in vitro

Taken together, the above results showing the *in vivo* clathrin uncoating defect and *in vitro* interaction between Hsc70 and α-synuclein suggest that when in excess, α-synuclein may interact with Hsc70 and alter its uncoating function at synapses. Amongst the possible mechanisms, α-synuclein could interfere with Hsc70 uncoating activity and/or alter Hsc70 availability at synapses. We began testing these possibilities by examining whether α-synuclein affects the ability of Hsc70 to promote clathrin cage disassembly *in vitro* using a dynamic light scattering assay as described in our previous studies (Jiang et al., 2005; Schuermann et al., 2008; Morgan et al., 2013). In one set of experiments we used empty clathrin cages assembled *in vitro* (Fig. 4A). In a typical reaction, Hsc70 binds the clathrin cages and subsequently uncoats clathrin, as indicated by a transient increase in light scattering followed by an exponential decrease in scattering intensity as the clathrin is disassembled (Fig. 4A, blue trace). Even when α-synuclein was in excess of Hsc70, it had no effect on the disassembly of clathrin cages (Fig. 4A, red trace). It is well established that α-synuclein interacts *in vitro* with small vesicles containing anionic lipids and *in vivo* with synaptic vesicles, and such vesicle interactions promote proper folding of the N-terminal α-helix (Maroteaux et al., 1988; Davidson et al., 1998; Fortin et al., 2005; Burre et al., 2015). Thus, because α-synuclein effects on uncoating may require membrane binding, we repeated the *in vitro* clathrin uncoating experiments using CCVs purified from bovine brains, which contained the underlying endocytic vesicle membranes (Fig. 3B). Hsc70 also bound and disassembled purified CCVs, although the decay in light scattering was less pronounced due to the presence of the vesicular membranes in the reaction (Fig. 4B, blue trace). Similar to the effects on clathrin baskets, α-synuclein also had no clear effect on Hsc70-mediated uncoating of purified CCVs (Fig. 4B, red trace). We thus conclude that α-synuclein does not affect the kinetics of clathrin uncoating *in vitro.*

**FIGURE 4.**
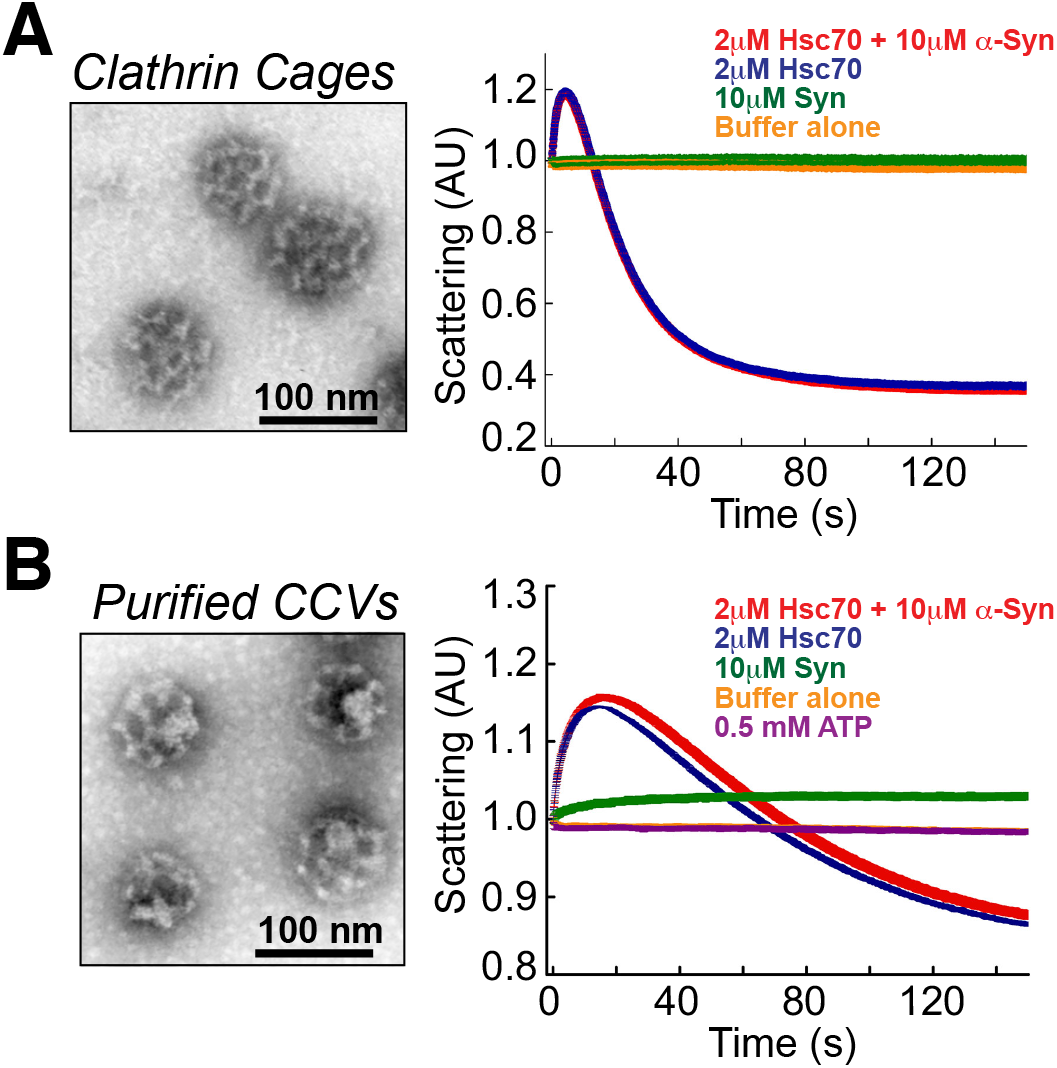
α-Synuclein does not affect Hsc70-mediated clathrin disassembly *in vitro.* **A.** (Left) Clathrin cages were assembled *in vitro* using recombinant clathrin heavy chain and auxilin, as described in (Sousa et al., 2016). (Right) Dynamic light scattering assay showed that addition of 2 μM Hsc70 exponentially reduced the light scattering intensity as clathrin cages were disassembled (blue trace). Addition of 10 μM α-synuclein did not alter the rate of Hsc70-mediated clathrin disassembly (red trace). No change in light scattering was observed in baseline control measurements after addition of buffer (yellow trace) or 10 μM α-synuclein alone (green trace). **B.** (Left) CCVs, which contain the underlying endocytic vesicle membranes, were freshly purified from bovine brains. (Right) Similar to the results with clathrin cages, introduction of 10 μM α-synuclein did not significantly affect the dynamics of Hsc70-mediated CCV uncoating (blue vs. red trace). α-Synuclein alone slightly increased light scattering (green trace), which is likely due to its binding to the endocytic vesicles.

### Excess α-synuclein inhibits Hsc70 availability at synapses

We next examined whether excess α-synuclein affects the localization of Hsc70 at the lamprey giant RS synapses. To do so, we performed whole mount immunostaining in the lamprey spinal cord for Hsc70 and SV2, a synapse marker, and then imaged the Hsc70 in the vicinity of the giant synapses. The giant RS synapses are large *en passant* synapses (1-2 μm in diameter; 1000-2000 synaptic vesicles), which reside along the periphery of the giant RS axons (see Fig. 1B, inset). They comprise large synaptic vesicle clusters at the active zone, which are surrounded by a distinct actin-rich periactive zone where clathrin-mediated synaptic vesicle recycling occurs (Fig. 5A) (Shupliakov et al., 2002; Morgan et al., 2004; Bourne et al., 2006; Saheki and De Camilli, 2012). To detect Hsc70, we used an Hsc70 antibody (Aviva ARP48445 rabbit polyclonal) that detects a single 70 kDa band in both lamprey and rat brain lysates by Western blotting (Fig. 5B). After immunostaining the spinal cords, the giant RS synapses were first identified using a monoclonal SV2 antibody, which labels presynapses in all vertebrates tested, including lampreys (Fig. 5C) (Buckley and Kelly, 1985; Busch et al., 2014). The synapse-associated Hsc70 was subsequently evaluated. At unstimulated control synapses, Hsc70 levels were low and diffuse, and distinct Hsc70 puncta were fairly sparse (Fig. 5C, top row). In contrast, stimulation (using high K^+^ ringer) induced an increase in Hsc70 puncta within or adjacent to the synaptic vesicle clusters, indicating enhanced Hsc70 availability at synapses (Fig. 5C, second row). After introducing excess human α-synuclein into the axons via microinjection (~7-16 μM as in the EM experiments), the stimulation-dependent increase in Hsc70 at presynapses was no longer observed (Fig. 5D). Next, we quantified all of the Hsc70 puncta within a 1-μm radius of the synaptic vesicle cluster, representing those within the endocytic periactive zone (see Fig. 5A). In unstimulated control axons, less than 20% of synapses per axon had clearly defined Hsc70 puncta associated with them (Fig. 5E). Stimulation induced a significant 3-fold increase in the percentage of synapses with Hsc70 puncta, and this did not occur after introduction of excess α-synuclein into the axons (Fig. 5E) (Unstim Control: 16.82 ± 3.76%, n = 84 synapses, 9 axons; Stim Control: 54.20 ± 5.28%, = 126 synapses, 12 axons; Unstim α-Syn: 18.09 ± 2.60 %, n = 65 synapses, 4 axons; Stim α-Syn: 27.98 ± 7.33 %, = 76 synapses, 6 axons; ANOVA p<0.0001; Tukey’s post hoc). We also quantified the average number of Hsc70 puncta per synapse under the same conditions and obtained similar results. At control synapses, the average number of Hsc70 puncta per synapse increased upon stimulation, but this did not occur after α-synuclein treatment (Fig. 5F) (Unstim Control: 0.19 ± 0.05 puncta, n= 84 synapses, 9 axons; Stim Control: 0.74 ± 0.09 puncta, n=126 synapses, 12 axons; Unstim α-Syn: 0.23 ± 0.02 puncta, n=65 synapses, 4 axons; Stim α-Syn: 0.37 ± 0.13 puncta, n=76 synapses, 6 axons; ANOVA p=0.0002, Tukey’s post hoc). In an independent set of experiments, using action potential stimulation (20 Hz, 5 min, as in the EM experiments) and another Hsc70 antibody (Enzo SPA815 rat polyclonal), we observed similar results. α-Synuclein eliminated the stimulation dependent recruitment of Hsc70 to synapses (Fig. G-H) (% Synapses w/Hsc70 Puncta: Unstim Control: 19.52 ± 5.37%, n = 34 synapses, 3 axons; Stim Control: 69.90 ± 5.15%, = 47 synapses, 3 axons; Unstim α-Syn: 32.50 ± 11.09 %, n = 31 synapses, 4 axons; Stim α-Syn: 33.33 ± 3.33 %, = 30 synapses, 3 axons; ANOVA p<0.01; Tukey’s post hoc) (#Hsc70 Puncta/Synapse: Unstim Control: 0.33 ± 0.12%, n = 34 synapses, 3 axons; Stim Control: 1.01 ± 0.21%, = 47 synapses, 3 axons; Unstim α-Syn: 0.50 ± 0.19 %, n = 31 synapses, 4 axons; Stim α-Syn: 0.47 ± 0.03 %, = 30 synapses, 3 axons; ANOVA p=0.08; Tukey’s post hoc). Thus, α-synuclein sequesters Hsc70 *in vivo* and reduces its availability at stimulated synapses, suggesting a possible mechanism by which α-synuclein causes CCV uncoating defects during synaptic vesicle recycling.

**FIGURE 5.**
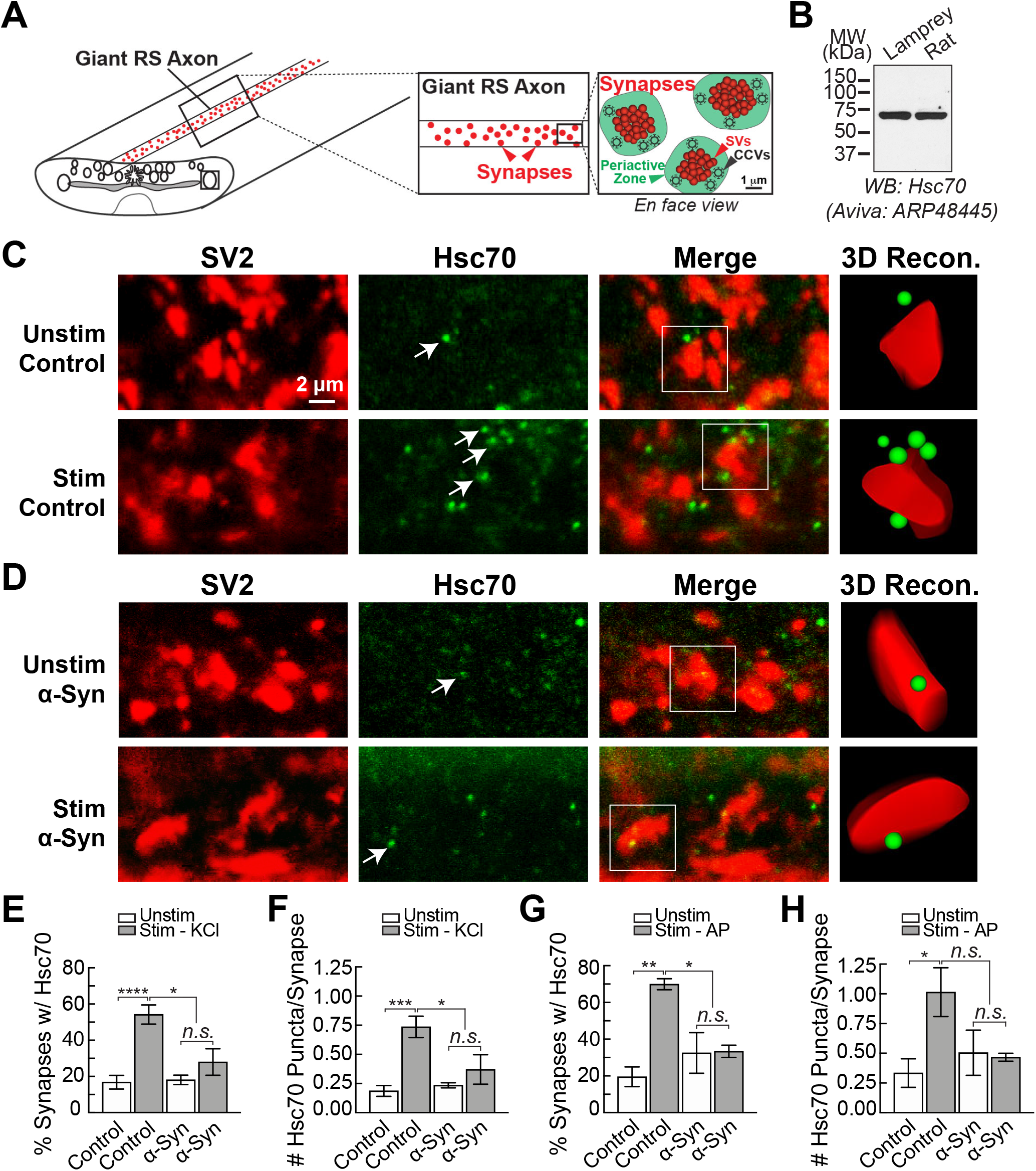
Excess α-synuclein inhibits Hsc70 availability at lamprey synapses. **A.** Diagram showing the basic organization of the giant RS synapses within lamprey spinal cord. The large synaptic vesicle (SV) clusters (red) are surrounded by a distinct periactive zone where clathrin-mediated endocytosis occurs (green). **B.** By Western blotting, the Hsc70 antibody used for these experiments (Aviva ARP48445) specifically recognized a single band at 70 kDa in both lamprey CNS and rat brain lysates, consistent with the expected molecular weight of Hsc70. **C.** Confocal images showing clusters of giant synapses immunostained for the synaptic vesicle-associated protein SV2 (red) and Hsc70 (green). Compared to unstimulated conditions, stimulation with high K^+^ increased Hsc70 availability at synapses, as evidenced by an increase in the number of visible Hsc70 puncta (arrows). **D.** In contrast, α-synuclein inhibited the stimulation-dependent increase in Hsc70 at synapses. **E-F.** Data showing the percentage of synapses (per axon) with associated Hsc70 puncta, as well as the average number of Hsc70 puncta per synapse. Bars represent mean ± SEM from n=65-126 synapses, 4-12 axons. * p<0.05, ** p<0.01, *** p<0.001, **** p<0.0001, and n.s.= not significant by ANOVA. **G-H.** Similar results were obtained using action potential stimulation (20 Hz, 5 min) and a different Hsc70 antibody for detection (Enzo Life Sciences SPA815). Bars represent mean ± SEM from n=30-47 synapses, 3-4 axons. * p<0.05, ** p<0.01, and n.s.= not significant by ANOVA.

### Increasing Hsc70 ameliorates the a-synuclein-induced vesicle trafficking defects

The above results suggest that reduced levels of Hsc70 at synapses –and consequently its function - underlie the α-synuclein-induced defects in CCV uncoating and synaptic vesicle recycling. If so, then increasing Hsc70 levels is expected to reverse the synaptic defects. To test this, we co-injected recombinant bovine Hsc70 along with human α-synuclein and examined the effects at lamprey synapses. As before, stimulated control synapses exhibited large vesicle clusters, shallow membrane evaginations, and few CCPs and CCVs (Fig. 6A,C). Compared to synapses treated with α-synuclein alone, which exhibited dramatic CCV uncoating and synaptic vesicle recycling defects (Fig. 1), those treated with α-synuclein and bovine Hsc70 together appeared relatively normal with large synaptic vesicle clusters, as well as few cisternae, CCPs, and CCVs (Fig. 6B,D). 3D reconstructions revealed the extent to which Hsc70 ameliorated the α-synuclein-associated synaptic vesicle trafficking defects (Fig. 6C-D). After introducing exogenous bovine Hsc70 along with human α-synuclein, the CCPs and CCVs were now sparse and located near the plasma membrane, similar to their distribution at control synapses (Fig. 6C-D, insets). Quantitative analyses revealed no difference in the number of synaptic vesicles at synapses co-treated with Hsc70+α-synuclein versus controls (Fig. 6E) (Control: 105 ± 11 SVs/section; Hsc70+α-Syn: 113 ± 13 SVs/section; n=22-30 synapses, 2 axons; Students t-Test; p=0.644). The synaptic vesicles were of similar size (diameter) in both conditions (Control: 54.6 ± 0.6 nm, n = 200 SVs, 10 synapses; Hsc70 + α-Syn: 52.9 ± 0.8 nm; n=200 SVs, 10 synapses; Student’s t-Test; p=0.07). Similarly, the number and size of cisternae were unchanged (Fig. 6G-H) (#Cisternae - Control: 2.7 ± 0.3 cisternae/section; Hsc70+α-Syn: 3.3 ± 0.4 cisternae/section; n=20-28 synapses; Students t-Test; p=0.26) (Size Cisternae - Control: 0.42 ± 0.02 μm; Hsc70+α-Syn: 0.45 ± 0.02 μm; n=53-91 cisternae, 20-28 synapses; Students t-Test; p=0.34). Hsc70 additionally restored the total number of clathrin coated structures (CCPs + CCVs) to control levels (Fig. 6I) (Control: 2.1 ± 0.3 coats/section; Hsc70+α-Syn: 2.4 ± 0.2 coats/section; n=20-28 synapses; Students t-Test; p = 0.35). Furthermore, there was a complete rescue of the CCV uncoating defect (Fig. 6J) (*Stage 1* - Control: 0.05 ± 0.05 CCPs; Hsc70+α-Syn: 0.07 ± 0.05 CCPs; *Stage 2–* Control: 0.40 ± 0.11 CCPs; Hsc70+α-Syn: 0.36 ± 0.12 CCPs; *Stage 3* - Control 1.25 ± 0.22 CCPs, Hsc70+α-Syn: 1.25 ± 0.20 CCPs; *Stage 4–* Control: 0.35 ± 0.11 CCVs, Hsc70+α-Syn: 0.68 ± 0.14 CCVs; n=20-28 synapses; ANOVA p=3.9×10^−12^, Tukey’s post hoc). Only the plasma membrane evaginations remained larger after co-injection of Hsc70 and α-synuclein, indicating some persistent effects on the plasma membrane (Fig. 6F) (Control: 1.95 ± 0.15 μm/section; Hsc70+α-Syn: 2.53 ± 0.20 μm; n=22-30 synapses; Students t-Test; p=0.04). The total membrane analysis further corroborated that synapses co-treated with Hsc70 and α-synuclein were similar to controls (Fig. 6K) (Control 1.4 ± 0.1 μm^2^; α-Syn: 1.5 ± 0.1 μm^2^; n=20-28 synapses; Student’s T-test; p=0.70). Thus, increasing exogenous Hsc70 levels largely reversed the CCV uncoating and synaptic vesicle recycling defects caused by α-synuclein.

**FIGURE 6.**
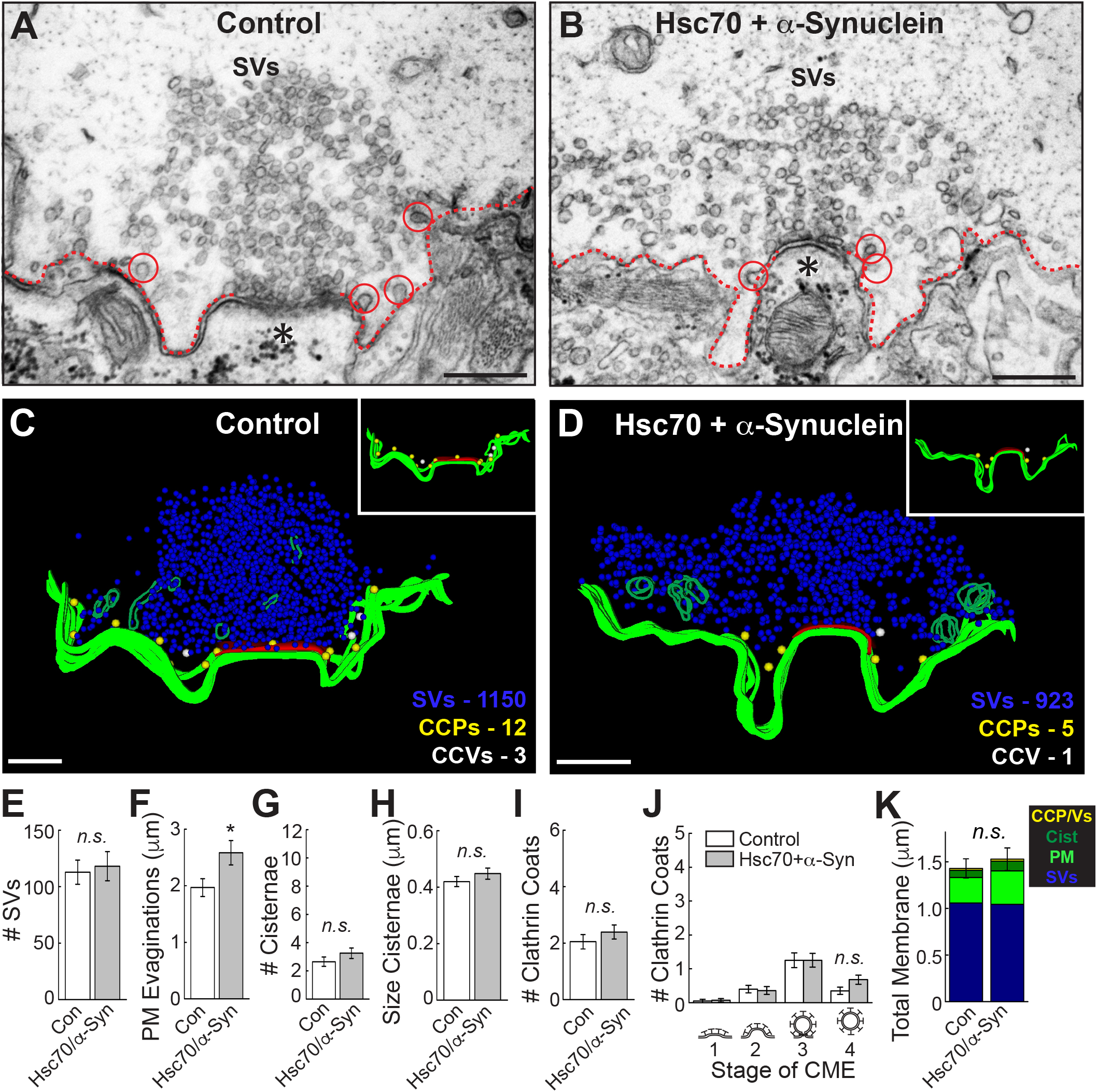
Increasing exogenous Hsc70 reverses the α-synuclein-induced synaptic defects. **A-B.** Unlike synapses treated with α-synuclein alone (see Fig. 1), those co-treated with Hsc70 and α-synuclein appear similar to control synapses, with large SV clusters and few cisternae or CCP/Vs (circles). **C-D.** 3D reconstructions reveal that synapses treated with Hsc70 and α-synuclein appear normal. Insets show the distributions of CCPs (yellow spheres) and CCVs (white spheres), which are sparse and clustered around the plasma membrane (green slabs). Active zone is shown in red. **E-K.** The CCV uncoating and vesicle recycling defects caused by α-synuclein were nearly ameliorated by co-injection of Hsc70 as evidenced by normal numbers of SVs, cisternae, and CCPs/CCVs. Only the PM evaginations were larger. Notably, there was no longer any difference in the number of free CCVs after co-injection of Hsc70 + α-synuclein, indicating a reversal of the uncoating defects (panel J, stage 4). Bars represent mean ± SEM (per section, per synapse) from n=22-30 synapses, 2 axons/condition. Asterisk indicates significance (p<0.05), and *n.s.* = not significant (p>0.05) by Student’s t-test (E-I, K) or ANOVA (J).

## DISCUSSION

While several studies have focused on the physiologic roles of α-synuclein at synapses, precise pathologic effects of α-synuclein overexpression are less clear. Data presented here identify loss of Hsc70 availability at synapses, and consequently its function, as one mechanism by which excess α-synuclein causes synaptic vesicle trafficking defects. Specifically, the sequestration of Hsc70 leads to an impairment of CCV uncoating at synapses, which consequently inhibits synaptic vesicle recycling (Fig. 7A). How might this work? A plausible explanation is that inhibiting CCV uncoating may trap clathrin and/or other limited coat proteins such as AP180 and AP2 within CCVs, making them unavailable for initiating subsequent rounds of endocytosis and leading to membrane expansion (Fig. 7A) (Morgan et al., 1999; Morgan et al., 2013). Membrane expansion may also be due to pleiotropic effects of excess α-synuclein on early stages of endocytosis, which is in line with our current understanding of its normal function (Vargas et al., 2014). Similar impairments of CCV uncoating have also been observed at squid and mammalian synapses after directly perturbing Hsc70 recruitment to CCVs (Morgan et al., 2001; Leshchyns’ka et al., 2006). As with direct Hsc70 perturbations (Morgan et al., 2001), excess α-synuclein also induced a loss of synaptic vesicles and expansion of the plasma membrane, indicating effects on vesicle endocytosis (Fig. 1) (Busch et al., 2014; Medeiros et al., 2017).

**FIGURE 7.**
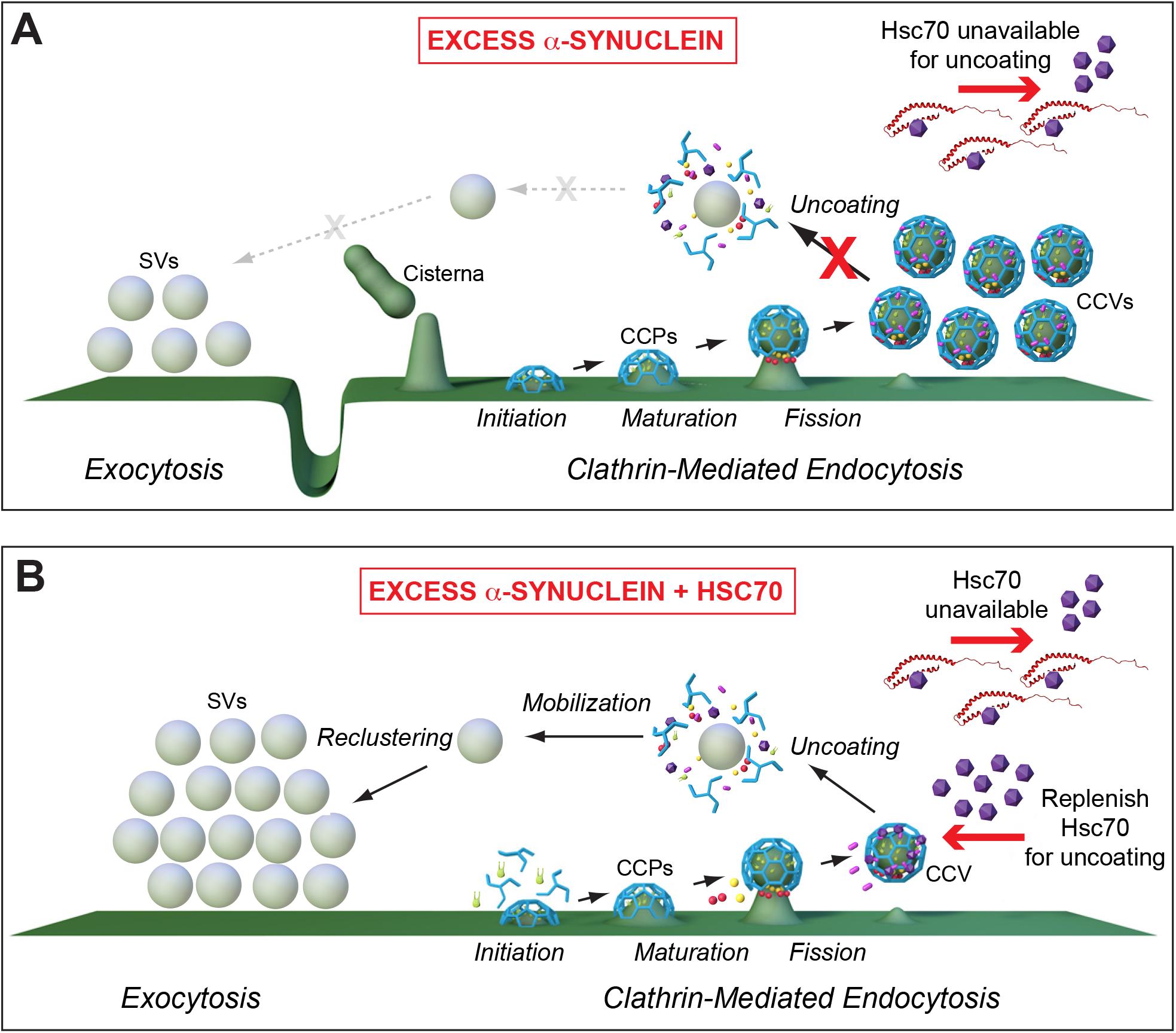
Model for α-synuclein-induced endocytic defects and amelioration by Hsc70. **A.** In the presence of excess α-synuclein, endogenous Hsc70 becomes depleted at synapses, leading to impaired CCV uncoating and an inhibition of synaptic vesicle recycling. **B.** Addition of exogenous Hsc70 restores CCV uncoating and leads to more normal synaptic vesicle recycling. (Graphics by Jack Cook, Woods Hole Oceanographic Institution).

By this working model, replenishing Hsc70 at synapses should have the effect of restoring CCV uncoating, thereby freeing clathrin and other coat proteins to recycle synaptic vesicles more efficiently. Indeed, when exogenous bovine Hsc70 was introduced along with human α-synuclein, the CCV uncoating defects were completely rescued, and the synaptic vesicle clusters were restored, indicating an improvement in vesicle recycling (Fig. 7B). After co-injection of Hsc70 and α-synuclein, the only morphological deficit remaining was an enlargement of the plasma membrane evaginations (Fig. 6). One possible explanation is that there is a pool of α-synuclein bound tightly to the membrane at the cell surface, which doesn’t release even in the presence of Hsc70 chaperone protein and therefore continues to slow membrane recycling from the plasma membrane. Another plausible explanation is that the excess Hsc70 sequesters clathrin, either by direct binding or by sequestration of other proteins required to release clathrin from endogenous Hsc70, which would also slow vesicle endocytosis. We previously showed that acute perturbations that result in persistent sequestration of clathrin by Hsc70 induced a similar phenotype: the appearance of plasma membrane evaginations consistent with impaired endocytosis, but without the appearance of CCVs (Morgan et al., 2013).

Our results corroborate and extend previous reports of an interaction between α-synuclein and Hsc70 (Pemberton et al., 2011; Redeker et al., 2012). Hsc70 binds both soluble and fibrillar α-synuclein in the absence of nucleotides and inhibits fibril formation *in vitro,* leading to increased cell viability (Pemberton et al., 2011). Cross-linking studies using soluble α-synuclein and Hsc70 indicated two discrete regions in the N-terminal domain of α-synuclein that bind to multiple sites within the client binding domain of Hsc70, including several in the C-terminal region (Redeker et al., 2012). Similar binding sites were observed between α-synuclein and the yeast Hsc70 ortholog, Ssa1p, indicating conservation of the interaction (Redeker et al., 2012). Our results further demonstrate conservation across synuclein orthologs (i.e. lamprey and human), which is due to the high degree of amino acid identity shared between their N-terminal domains (Busch and Morgan, 2012). Our data also corroborated the NTD of α-synuclein as the region that binds to Hsc70. Finally, we show that the C-terminal region of Hsc70 is critical for mediating and/or stabilizing the interaction with α-synuclein. The failure of Hsc70ΔC to bind α-synuclein is consistent with observations that the region deleted in Hsc70ΔC includes residues that form part of the interface with α-synuclein (Redeker et al., 2012). In addition, the deletion in Hsc70ΔC destabilizes the terminal helix of Hsc70 SBD, resulting in unwinding of this helix and cis-binding of its unfolded segment in the client binding site of nucleotide free Hsc70 (Morshauser et al., 1999; Jiang et al., 2005). The client binding site in Hsc70 also contributes to the interface with α-synuclein (Redeker et al., 2012), and so cis-binding of this unfolded segment may compete with α-synuclein binding. Going forward, it will be important to identify the most critical residues mediating the α-synuclein/Hsc70 interaction in order to facilitate the design of reagents that disrupt this interaction, which might have therapeutic value for treatment of multiple diseases associated with α-synuclein and/or clathrin uncoating defects.

Hsc70 was previously identified as a potential target in other α-synuclein-related contexts. In human brains affected by PD, DLB and related synucleinopathies, Hsc70 and other chaperone proteins are detected in high levels along with α-synuclein (Auluck et al., 2002; Uryu et al., 2006). The co-occurrence of Hsc70 and α-synuclein in Lewy bodies suggests an increased association of these proteins in disease states, which could deplete Hsc70 and impair its function in other cellular compartments. In support of this idea, α-synuclein aggregation, fibrillation, trans-synaptic propagation, and neurotoxicity are reduced in the presence of Hsc70 or Hsp70 (Luk et al., 2008; Danzer et al., 2011; Pemberton et al., 2011). Furthermore, overexpression of Hsp70 reduced α-synuclein aggregation and neuronal loss in *Drosophila* and mouse models of α-synuclein toxicity (Auluck et al., 2002; Klucken et al., 2004). Thus, Hsc/p70-based reagents that can reverse misfolding and/or restore normal protein function have been suggested as potential therapeutic agents to reduce neurodegeneration caused by α-synuclein (Dimant et al., 2012; Ebrahimi-Fakhari et al., 2013; Shorter, 2016). Our data extend the possible applications of such agents by showing that replenishing Hsc70 function is also a viable strategy for improving the synaptic defects observed in α-synuclein-related disorders.

Importantly, our work also provides new insights into the cellular mechanisms for α-synuclein and Hsc70 involvement in the neurodegenerative process, specifically related to the impacts on CME. Since CME is also involved in vesicle endocytosis from the plasma membrane (e.g. receptor internalization) and intracellular vesicle trafficking (e.g. trans-Golgi to endosomal transport), these experiments also have wider implications for impacts on membrane trafficking events throughout the entire neuron. In addition, our work also highlights the need for the type of studies described here, in which the mechanism of phenotypic rescue is characterized in depth, because of the potential complexities of deploying a therapeutic approach that involves increasing the activity of a protein like Hsc70, which is involved in many cellular processes. We find that while Hsc70 fully rescues the CCV uncoating defects induced by α-synuclein and restored the vesicle cluster, the membrane evaginations persisted, suggesting residual impairment of membrane endocytosis by α-synuclein. Alternatively, the enlarged plasma membrane evaginations may result not from a failure to rescue, but to a direct consequence of increased Hsc70 levels (Morgan et al., 2013). This suggests not only the need to titrate such therapeutic approaches, but to understand the mechanisms of phenotypic rescue so that any complications can be anticipated.

The findings reported here contribute to a growing body of evidence that PD and other forms of Parkinsonism may be linked to deficits in the clathrin-mediated synaptic vesicle recycling (Saez-Atienzar and Singleton, 2017), and an increasing number of findings are pointing toward specific impairments in the clathrin uncoating process. For example, many studies have now identified mutations or truncations in *DNAJ*, the gene that encodes for auxilin (the Hsc70 cochaperone at synapses) in patients with juvenile Parkinsonism or early-onset Parkinson’s disease (Edvardson et al., 2012; Koroglu et al., 2013; Elsayed et al., 2016; Olgiati et al., 2016). In addition, auxilin was recently identified as a target of the PD-linked LRRK2 mutant R1441C, which led to impaired endocytosis and clathrin uncoating defects in patient-derived dopaminergic neurons (Nguyen and Krainc, 2018). Furthermore, GWAS studies have identified genetic variations and increased expression of GAK, the ubiquitously expressed Hsc70 co-chaperone, amongst the top risk factors for familial PD across multiple populations worldwide (Pankratz et al., 2009; Tseng et al., 2013; Nagle et al., 2016). Several mutations in synaptojanin 1 have also been linked to early onset Parkinsonism (Krebs et al., 2013; Chen et al., 2015; Ben Romdhan et al., 2018). Transgenic mice carrying one of the synaptojanin I mutations (R258Q) in the Sac domain accumulated CCVs within their synapses and exhibited impaired synaptic vesicle recycling, as well as motor deficits and increased death (Cao et al., 2017). Thus, a growing body of evidence from both animal models and human genetics indicates that defects in CME, and specifically in CCV uncoating, are at least susceptibility factors in PD and Parkinsonism, if not causal factors. Thus, strategies that ensure proper CCV uncoating, for example by increasing Hsc70 availability or function, may hold promise for improving synaptic function and reducing neurodegeneration in PD and other α-synuclein-associated diseases.

## ACKNOWLEDGEMENTS

This work was supported by grants from the NIH NINDS/NIA R01NS078165 (to JRM), NIH/NIGMS R01GM118933 (to EML and RS). We thank Paul Oliphint and Rylie Walsh for technical support; and Dr. Julia George for the gift of α-synuclein constructs and early discussions. Thanks to Louie Kerr and Kasia Hammar from the MBL Central Microscopy Facility for providing EM support.

